# The impact of subunit type, alternative splicing, and auxiliary proteins on AMPA receptor trafficking

**DOI:** 10.1101/2024.12.18.629280

**Authors:** Tyler Couch, Tyler W. McCullock, David M. MacLean

## Abstract

AMPA receptors underlie fast excitatory synaptic transmission in the mammalian nervous system and are critical for the expression of synaptic plasticity. Four genes encode the AMPA subunits, each subject to RNA editing and alternative splicing at multiple positions. In addition, each tetrameric AMPA receptor can harbor up to four auxiliary proteins, of which there are multiple types. Subunit type, alternative splicing, and auxiliary proteins are all known to affect AMPA receptor gating and trafficking. However, determining which factors dominate AMPA receptor trafficking requires high throughput assessment of trafficking across multiple conditions. Here, we deploy two such methods to assess the relative contribution of AMPA subunit type (GluA1 versus GluA2), alternative splicing (flip versus flop), and various transmembrane AMPA receptor regulatory proteins (TARPs) to AMPA receptor trafficking. We find that subunit type is the most important factor, with GluA2 showing much better surface expression than GluA1, and alternative splicing plays a secondary role, with flip subunits consistently outperforming flop variants in surface expression across all conditions. Type 1 TARPs (γ2-4 and γ8) enhance surface trafficking, while Type 2 TARPs (γ5 and γ7) reduced surface expression, although we could not detect differences within each type. These data will be a helpful resource in comparing surface expression across a variety of AMPA receptor compositions. Our assays will also enable high throughput assessment of novel disease-associated mutations, chimeras, and auxiliary and chaperone proteins.

## Introduction

AMPA receptors mediate the majority of fast excitatory transmission in the mammalian central nervous system^1^. These ionotropic glutamate receptors are tetrameric assemblies of various subunits (GluA1-4) that can form homomeric or heteromeric complexes. Dynamic regulation of synaptic AMPA receptor number and type is a powerful method of controlling synaptic strength in response to activity and during development^1–3^. Regulating synaptic AMPA receptor content involves shifting the balance between endo- and exocytosis as well as lateral diffusion from extra-synaptic spaces^3–5^. In addition, local synthesis of new AMPA subunits in the dendritic spine can contribute additional AMPA receptors as needed^6,7^. Supplying new AMPA receptors to synapses through any of these routes ultimately depends on efficient endoplasmic reticulum synthesis and exit^8^. Multiple factors contribute to the efficiency of AMPA receptor synthesis or biogenesis, including the type of subunit (GluA1 versus GluA2, 3 or 4), RNA editing^9,10^, alternative splicing^11,12^, and the presence of chaperone^13^ and auxiliary proteins^1,14–16^.

Native AMPA receptors assemble with a myriad of auxiliary proteins^1,17^. These include the cornichon proteins^18^, cys-knot associated proteins (CKAMP)^19^, GSG1L^20,21^, and the transmembrane AMPA receptor regulatory proteins (TARPs)^14,22^. TARPs generally act as gain-of-function auxiliary proteins, increasing agonist efficacy and potency^23^, attenuating polyamine block^24^, slowing desensitization, and accelerating recovery^22,25^ (with some exceptions^26^). Most TARPs also promote the surface expression of AMPA receptors^22,27–29^ and their differential retention in synaptic versus extra-synaptic spaces^30^. Prior work has examined the impact and/or mechanism of specific TARPs such as γ2 (aka stargazin)^22,27,29^ or γ8, or compared measured the impact of several TARPs on AMPA receptor surface^31^. But to our knowledge, there has been no systematic assessment of the impact of AMPA subunit type, splice variant, and TARP on surface expression.

Here, we dissect the relative contributions of subunit composition, splice variants, and TARP auxiliary proteins in AMPA trafficking and any potential interactions between these factors through two distinct approaches. First, we implement a flow cytometry method to examine GluA1, GuA2, and GluA1/2 surface expression alone or in the presence of TARPs with single cell resolution. Second, we develop a split luciferase assay to measure the fraction of AMPA receptors on the cell surface across a population. We systematically measure the plasma membrane levels of GluA1 and GluA2, both flip and flop isoforms. For each subunit and heteromeric assembly, we additionally evaluate the impact of the TARP family of proteins, γ1-8 (excluding γ6). We find that GluA2 traffics more efficiently than GluA1 and flip isoforms traffic better than flop^12^. We also find that type 1 TARPs (γ2-4 and 8) enhance plasma membrane expression, while type 2 TARPs (γ5 and γ7) impair surface trafficking. In general, all these effects are additive but some combinations produce synergistic effects.

## Materials and Methods

### Plasmids and cloning

The cDNA encoding mouse γ1 was a kind gift from Dr. Robert Dirksen. γ2 (aka stargazin), γ3, γ4, γ8, GluA1, and GluA2 (short form, UniProt [P19491]) were kind gifts from Drs. Bowie or Jayaraman. γ5 and γ7 were synthesized as gBlocks (Integrated DNA Technologies). All TARP and AMPA subunit cDNAs were subcloned into pcDNA3.1(+) by PCR amplification with high-fidelity Q5 polymerase (New England Biolabs) using primers with restriction sites in the overhangs. A consensus Kozak sequence (GCCACC) was also added to precede the intended start codon. The mNeonGreen coding sequence was inserted at the carboxy terminus of GluA1 and GluA2 using NEBuilder HiFi DNA Assembly Master Mix from PCR-derived fragments (New England Biolabs). For flow cytometry, HA epitope tags were inserted following the signal sequences for GluA1 and GluA2 using Q5 mutagenesis/KLD enzyme mix (New England Biolabs). The sequence corresponding to the flip isoform was converted into the corresponding flop sequence using DNA fragments generated by polymerase chain assembly (GluA1) or a gBlock (GluA2) and NEBuilder HiFi DNA Assembly Master Mix. For luciferase-based surface trafficking, the signal peptides were replaced with that of the hemagglutinin signal peptide (HAsp)^32^, followed by an HA epitope, a GS linker, the 11-amino acid HiBiT tag^33^ and a GSTG linker before the start of the mature AMPA receptor sequence. All cDNA constructs were verified by Sanger sequencing (Eurofins Genomics) or Oxford Nanopore sequencing (Plasmidsaurus).

### Cell culture and transfection

FreeStyle™ 293-F (HEK293F) cells were purchased from ThermoFisher Scientific and maintained in FreeStyle™ 293 Expression Medium in a 125 ml tissue culture flask under constant shaking at 135 rpm according to the manufacturer’s instructions. For transfection, 5x10^5^ cells were seeded into each well of a 12-well plate and transfected with 1.5 µg of cDNA (0.75 µg TARP, 0.75 µg AMPA subunit) using a 3:1 mass ratio of PEI:cDNA. NBQX was added to a final concentration of 20 µM at the time of transfection to promote viability. The transfection plates were then returned to the shaking platform to keep cells in suspension.

### Flow cytometry

Two hours prior to flow cytometry analysis, cells were treated with 100 µM cycloheximide to inhibit protein synthesis and synchronize fluorescent protein maturation^34^. Cells were pipetted from their 12-well plate into 2 mL of FACS buffer (divalent-free DPBS containing 0.3% BSA and 1 mM Na_4_EDTA) and pelleted by centrifugation (200 x g, 5 minutes, 4 °C). The cell pellets were resuspended in 100 µL of staining mix (10% FBS in FACS buffer) and 2 µL of allophycocyanin (APC) anti-HA.11 [clone 16B12] (BioLegend). This amount of antibody was empirically tested to yield the highest stain index. Cells were stained on ice protected from light for 20 minutes before being washed with 4 mL of FACS buffer and recollected by centrifugation as before. Cells were resuspended in 100-200 µL of FACS buffer containing 0.2 ug/mL of DAPI (Cell Signaling Technologies) to assess viability.

Cells were run on an LSRII following instrument compensation collecting 50,000 events per condition. DAPI was excited with a 405 nm laser and detected between 425-475 nm, GFP with a 488 nm laser and detected between 505-525 nm, and APC with a 633 nm laser and detected between 650-670 nm. Single colored GFP samples were run after compensation to confirm fluorescence spillover was being compensated.

Data analysis was performed using FCS Express 7 Flow (De Novo Software). Single cells were gated for using forward scatter and side scatter, and only viable cells based on DAPI-exclusion were used for analysis (typically 25,000-30,000 events of the 50,000 collected). The Surface/Total GluA parameter was created using the parameter math function in FCS Express where the fluorescence intensity of the HA-APC channel was divided by the fluorescence intensity of the GFP channel for each individual cell.

### NanoBiT surface trafficking assay

HEK293F cells were added to a 12-well plate, 1 ml of 1x10^6^ cells/ml in FreeStyle media supplemented with 20 µM NBQX, and immediately transfected with 500 ng of AMPA receptor and 1000 ng of auxiliary protein or empty vector using PEI at a 3:1 PEI to DNA ratio. 20-24 hours post-transfection, 50 uL of cells from each transfection were transferred to a well of an opaque 96-well plate, supplemented with 50 uL of fresh detection reagent (see below), and incubated for at least 5 minutes. Luminescence measurements were conducted using FLUOstar plate reader (BMG) without a filter with readings once per minute and a 480 ms integration time. Following 10 minutes of acquisition, 11 uL of 4 mg/ml digitonin (in 90% PBS, 10 % DMSO) was manually mixed into each well. Luminescence was continuously measured for 30 minutes to monitor cell permeabilization by digitonin. The luminescence at the 10-minute time point (immediately before digitonin addition) and at the end of the 30-minute digitonin incubation were taken as the surface and total measurements, respectively. In all experiments, a non-transfected condition was included and used for background subtraction. For ‘wash’ experiments, cells were first collected, washed three times with PBS (supplemented with 20 µM NBQX), and re-suspended in 1 ml of PBS containing NBQX before 50 uL was taken for luminescence readings.

The detection reagent was a mix of in-house purified LgBiT protein and hydrolyzed Hikarazine 103, each diluted 1:200 in PBS. To purify LgBiT, a g-Block (IDT) encoding an *E.coli* optimized LgBiT coding sequence with N terminal hexa-histidine and TEV cleavage sequence was inserted into the pET28a(+) expression vector. A single colony of BL21(DE3) *E.coli* (NEB), transformed with this vector, was expanded to 2L with an OD_600_ of 0.6 using a constant temperature of 37°C. Expression was induced with 1 mM IPTG. Following overnight incubation at 18°C, bacteria were pelleted and stored at -80°C. Thawed bacteria were resuspended in purification buffer (20 mM sodium phosphate, 500 mM NaCl, pH 7.5) with all subsequent steps performed at 4°C, lysed by sonication, and incubated with 1 mg/ml Lysozyme for 1 hour. Lysates were cleared by centrifugation first at 6000g for 30 minutes, then 50,000g for 30 minutes, and finally filtered using 0.45 µM PVDF syringe filter. The cleared lysate was supplemented with 10 mM imidazole and flowed over a pre-equilibrated HisPur Ni-NTA Spin Column (Thermo Fisher). After 20 column volume washes in wash buffer (20 mM sodium phosphate, 500 mM NaCl, 25 mM imidazole pH 7.5), protein was eluted in 1 ml fractions using elution buffer (20 mM sodium phosphate, 500 mM NaCl, 250 uM imidazole, pH 7.5) and fractions analyzed using SDS-PAGE and luminescence. LgBiT-containing fractions were pooled, and buffer exchanged into storage buffer (20 mM sodium phosphate, 150 mM NaCl, 50% glycerol, pH 7.5) using a PD-10 desalting column (Cytiva). LgBiT protein was stored at -80°C for long-term or -20°C for short-term at concentrations comparable to commercial LgBiT protein, as estimated by SDS-PAGE and luminescence.

The o-acylated pro-luciferin Hikarazine 103 was purchased from Yves L. Janin (Museum National d’Histoire Naturelle, Paris)^35,36^ and converted to an active form by solubilizing 1 mg of Hikarazine 103 in 0.2 mL of DMSO, mixing 0.3 mL of acidic ethanol (0.1 mM HCl in 200 proof ethanol). The hydrolysis reaction was placed in a 50°C water bath for 2 hours, then transferred to -20°C for storage. Hydrolysis was performed in 1 mg batches as needed.

### Electrophysiology

Culture dishes were visualized with phase contrast on a Nikon Ti2 microscope using a 20x objective. Outside-out patches were excised using heat-polished, thick-walled borosilicate glass pipettes of 3 to 5 MΩ resistance. The pipette internal solution contained (in mM) 135 CsF, 33 CsOH, 11 EGTA, 10 HEPES, 2 MgCl_2_ and 1 CaCl_2_ (pH 7.4) and the external contained 150 NaCl, 10 HEPES, 1 MgCl_2_ and 1 CaCl_2_ and was supplemented with 10 mM monosodium glutamate as an agonist. All recordings were performed at room temperature with a holding potential of -60 mV using an Axopatch 200B amplifier (Molecular Devices) and Clampex 10 or 11. Data were acquired 50 kHz, filtered at 10 kHz with series resistance was routinely compensated by 90 to 95% where the peak amplitude exceeded 100 pA. Rapid perfusion was performed using home-built, double or triple-barrel application pipettes (Vitrocom), manufactured as described previously^37^. Translation of application pipettes was achieved using piezo actuators driven by voltage power supplies. The command voltages were generally low pass filtered (50-100 Hz, eight-pole Bessel).

### Statistics and Data Analysis

Log-log flow plots of the GFP versus APC signal were fit with a piecewise linear or segmented function with two phases or segments. Data where x < k, were fit with:

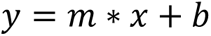

where *x* and *y* are the GFP and APC signal intensities, respectively, *m* is the slope, and *b* is the y-intercept signal intensity. The free parameter k is the ‘threshold’ and data where x > k were fit with:

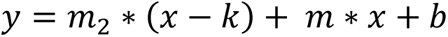

where *m_2_* is a second slope, reflecting the surface trafficking of receptors in excess of the threshold *k*.

Statistical comparisons between two samples were made using two-tailed heteroscedastic t-tests. Comparisons between many samples across conditions were done using two- and three-way ANOVAs, implemented in Prism (GraphPad). The Benjamini, Krieger and Yekutieli method of multiple comparison testing was used. P values less than 0.05 were statistically significant. A single transfection for any given condition, run on either flow cytometer or plate reader, was taken to be a biological replicate or n.

## Results

### Flow-based trafficking assay

To assay the plasma membrane levels of AMPA receptors in a quantitative, high throughput manner, we drew upon past flow-based strategies that measure the total receptor number using genetically encoded fluorescent proteins and the surface receptor number using an extracellular epitope and cell-impermeant labels (Figure 1A)^38–41^. We appended the bright and fast-maturing mNeonGreen^42^ (hereafter referred to as GFP) to the cytoplasmic tail of GluA1 and GluA2. An amino-terminal hemagglutinin (HA) tag between the signal peptide and the amino-terminal domain served as the extracellular epitope. These constructs were transfected into the FreeStyle 293-F suspension cell line, and the GFP and HA-APC emission intensities were measured using flow cytometry to assay total and surface receptor expression, respectively (gating scheme in Figure S1). However, each of these channels contains extraneous components. Since the GFP variant matures quickly (∼10 minutes^42^), the GFP channel reflects some fraction of mature GFP whose upstream AMPA subunit is still folding. We routinely added the protein synthesis inhibitor cyclohexamide before flow experiments to reduce this asynchrony. The HA-APC channel may contain signal from anti-HA antibodies trapped in dead or dying cells, leading to false positive surface signals. AMPA receptors, especially co-transfected with TARPs, can cause excitotoxicity and possibly cause even greater false positive surface signal (Figure S1B). Therefore, we use DAPI to exclude dead cells, minimizing the effect of anti-body trapping (Figure S1B). Example flow plots of the surface (APC) versus total (GFP) signals from GluA1 and GluA2 (both flip and flop variants) are shown in Figure 1B. As expected, GluA2 showed greater surface expression than GluA1 based on histograms of APC fluorescence intensity (Figure 1B, right side of each plot). GluA2 surface expression could be higher because GluA2 protein is simply produced more efficiently in recombinant systems. To control for this, we examined the surface-to-total signal ratio for each cell (e.g., single-event APC/GFP intensity), effectively normalizing the surface signal to total AMPA content in each cell. GluA2, both flip and flop, had higher surface/total ratios than GluA1, as seen in the right shifts of cumulative distribution plots towards the higher surface/total ratios (Figure 1C). Consistent with this, at the population level, GluA2 showed higher percentages of APC+ cells that were also GFP+ (Figure 1D, percentage APC+ of GluA2 flip: 66 ± 3%, n = 8; GluA2 flop: 51 ± 3%, n = 4; GluA1 flip: 40 ± 2%, n = 8, p = 8e-6 versus GluA2 flip; GluA1 flop: 25 ± 3%, n = 4, p = 0.0016 versus GluA2 flop). The flip variants of both GluA1 and GluA2 had stronger surface trafficking than their flop counterparts, also consistent with past work^12,43^.

**Figure 1.**
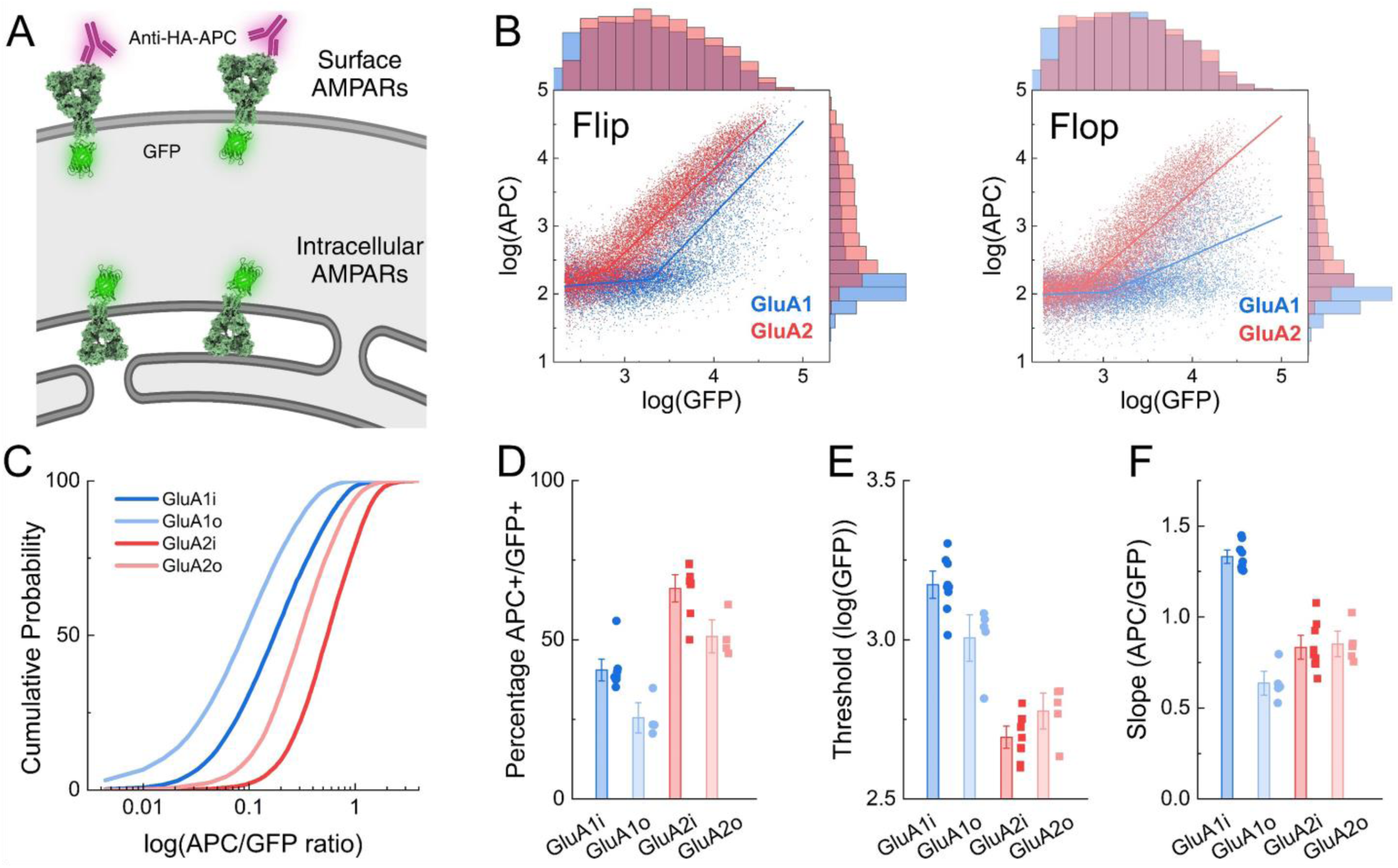
Intrinsic AMPA subunit and splice variant trafficking differences revealed by flow cytometry-based assay. **(A)** Schematic of flow-based assay where intracellular green fluorescence reports total AMPA receptors and extracellular APC fluorescence indicates the plasma membrane component. **(B)** Dot plots for flip *(left)* and flop *(right)* variants of GluA1 *(blue)* and GluA2 *(red)* showing the total AMPA receptors on the x axis (GFP channel) and surface AMPA receptors on the y axis (HA-APC) in log scale. Solid line is a piecewise linear fit. Upper histogram represents GFP counts while APC counts are shown on the righthand histogram. **(C)** Cumulative probability of single event APC over GFP ratios for GluA1 *(blue)* or GluA2 *(red)* subunits, both flip *(i, darker colors)* and flop *(o, lighter colors)* for the experiment in **B. (D-F)** Summary of the percentage of APC+ cells within the GFP+ population **(D),** threshold **(E)** and **(F)** slope of piecewise fit of the indicated constructs. Symbols are separate transfections and error bars are SEM.

A striking feature of the flow plots is a ‘hockey stick’ pattern. Phenotypically, the GFP signal intensity increases without much change to the APC signal until, at some point, both signals increase. Thus, AMPA receptors seem to accumulate intracellularly until reaching some threshold, at which point they emerge on the plasma membrane. However, this point of emergence is different for each subunit, with the GluA2 flip beginning to show detectable plasma membrane expression at much lower levels of total AMPA receptor (e.g. GFP signal) than the GluA1 flip (Figure 1B, left). A similar effect was observed with GluA1 and GluA2 flop. However, once these receptors began to appear on the surface, the slope of the APC/GFP relation was quite different (Figure 1B, right). We quantified this hockey stick pattern using a piecewise linear function (see Methods) that separates the data into phases with distinct slopes. In the ‘initial’ phase, AMPA receptors accumulate in cells but with little surface signal; hence, the slope is small. In the ‘later’ phase, AMPA receptors continue accumulating in cells but also appear on the surface; hence, the slope is steeper. We termed the transition between these two phases the “threshold”. The threshold for GluA2, both flip and flop, was lower than that of GluA1 (Figure 1E, threshold (log(GFP)) GluA2 flip: 2.69 ± 0.02, n = 8; GluA2 flop: 2.77 ± 0.04, n = 4; GluA1 flip: 3.17 ± 0.03, n = 8, p = 1e-9 versus GluA2 flip; GluA1 flop: 3.00 ± 0.05, n = 4, p = 0.006 versus GluA2 flop). Interestingly, the post-threshold slope of the APC/GFP signal also varied between subunits with the following rank order (steepest first): GluA1 flip, GluA2 flip = GluA2 flop, GluA1 flop (Figure 1F, slope (log(APC)/log(GFP)) GluA1 flip: 1.33 ± 0.02, n = 8; GluA2 flip: 0.83 ± 0.04, n = 8; GluA2 flop: 0.85 ± 0.05, n = 4, GluA1 flop: 0.64 ± 0.04, n = 4). Such differences may arise due to differential rates of folding, dimer and tetramer assembly, ER exit, processing, and the endo/exocytosis balance. The possible processes contributing to the threshold and slope measurements need further clarification (see Discussion). But empirically, GluA2 combined an intermediate slope with a lower threshold, while GluA1, both flip and flop, had a higher threshold (Figure 1E and F). Interestingly, GluA1 flip had the steepest slope while GluA1 flop had the shallowest (Figure 1F). Since AMPA auxiliary or chaperone proteins found in neurons are absent or present at low levels in HEK cells^13,44^, these observed differences arise from the intrinsic folding, multimerization, and trafficking of the AMPA receptor.

### TARP effects on AMPA surface expression as measured by flow cytometry

Next, we assessed the impact of TARP auxiliary proteins on the trafficking of GluA1 flip. TARPs are divided into type 1 and 2, with type 1 being further subdivided into type 1a and 1b^1^. To systematically assess how each of the TARP types impacts GluA1 trafficking, surface, and total GluA1 flip were measured in cells co-transfected with either empty vector (EV) or each of the TARPs. We included γ1 as an additional negative control as it bears sequence homology with TARPs but has been reported not to modulate AMPA currents^14^. Figure 2a shows an example of flow plots of GluA1 flip plus empty vector, type 1a TARP γ2 or type 2 TARP γ5. As expected, γ2 increased theslope of the APC/GFP ratio. Interestingly, γ5 co-transfection reduced or shallowed the slope. We found a similar pattern across all TARPs where type 1a and 1b TARPs enhanced the surface trafficking of GluA1 flip, producing right shifts in the cumulative probability of surface/total ratios compared to EV or γ1 co-transfection (Figure 2B). In contrast, the type 2 TARPs γ5 and γ7 left shifted the distributions, indicating impaired trafficking (Figure 2B). To further dissect TARP differences, we fit the single cell flow data (Figure 2C) to measure threshold and slope. No statistical effect was observed on the threshold of GluA1 flip for any TARP (Figure 2D, 1-way ANOVA F(7,16) = 0.7973, p = 0.6). However, both type 1a and 1b TARPs did increase the slope of GluA1 flip trafficking, while type 2 TARPs reduced the slope (Figure 2D, slope (log(APC)/log(GFP)) GluA1 flip with EV: 1.30 ± 0.06, n = 3; with Type 1 TARPs γ2-4, γ8 range from: 1.54 ± 0.03 to 1.63 ± 0.03, n = 3, p versus EV all less than 0.0001; with γ5: 0.94 ± 0.04, n = 3, p < 0.001 versus EV; with γ7: 0.67 ± 0.03, n = 3, p < 0.001 versus EV).

**Figure 2.**
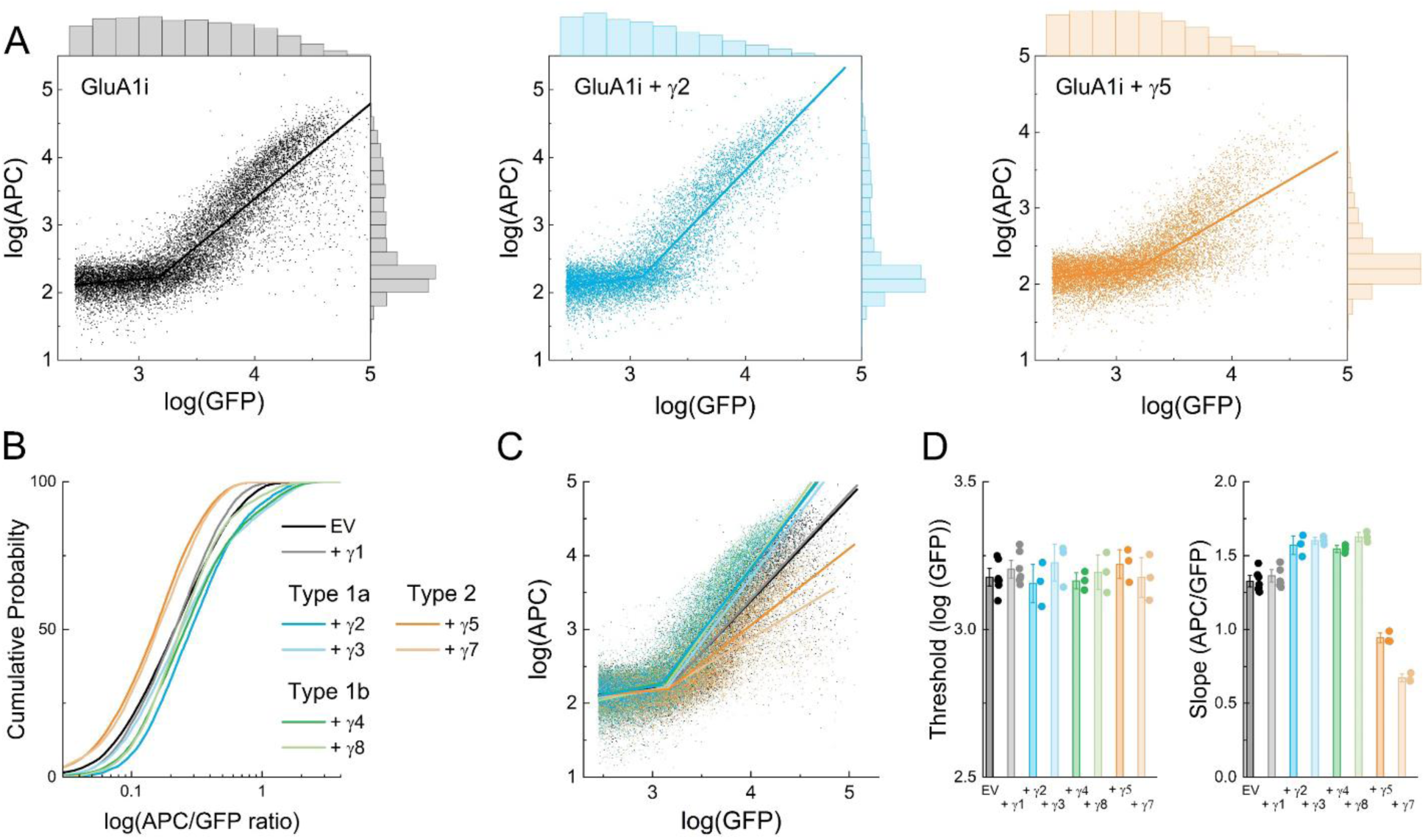
GluA1 surface trafficking is enhanced by Type la and 1b but reduced by Type 2 TARPs. **(A)** Dot plots for GluA1 flip either alone *(left, black),* or co-traπsfected with Type 1a TARP γ2 *(middle, blue)* or Type 2 TARP γ5 *(right, orange).* A solid line is a piecewise linear fit (see Methods). The upper histogram represents GFP counts, while APC counts are shown on the righthand histogram. (B) Cumulative probability of single event APC over GFP ratios for GluA1 flip alone *(black)* or with the indicated gamma subunit. (C) Dot plots for GluA1 flip alone *(black)* or with the stated gamma subunit sharing the color scheme of **B. (D)** Summary threshold *(left)* and slope *(right)* from linear piecewise fit for each gamma subunit with GluA1 flip. Symbols are separate transfections and error bars are SEM.

To determine if the impact of TARPs on AMPA trafficking depended on subunit identity or splice variants, we repeated this experiment with GluA1 or GluA2 (both flip and flop) as well as GluA1/GluA2 co-transfection composed solely of flip or flop subunits. The heatmaps in Figure 3A show the following general trends from these experiments: GluA2 had greater expression than GluA1 with heteromers falling in between, flip receptors had greater surface expression than flop, and type 1 TARPs increased surface trafficking while type 2 impaired it (Figure 3A). Representative single-cell surface/total ratios from one such experiment are shown in Figure S2. The percentage of jointly APC+ and GFP+ cells, single cell surface to total ratios as well as fit values of thresholds, and slopes from these experiments are all summarized in Figures S3 and S4 and Tables 1-4. To estimate the relative influence of subunit, splice variant, and TARP on these observable trafficking markers (%APC+ of GFP+ and surface/total ratio), we conducted a three-way ANOVA on the homomeric data with subunit identity (GluA1 versus GluA2), splice variant (flip versus flop), and TARP as factors. All of these factors showed significant effects on the observable trafficking markers. Specifically, subunit identity showed the strongest influence (%APC+ of GFP+: 60% of variance, F(1,64) = 823, p < 0.0001; APC/GFP ratio: 70% of variance, F(1,64) = 1120, p < 0.0001) followed by the flip/flop cassette (%APC+ of GFP+: 18% of variance, F(1,64) = 249, p < 0.0001; APC/GFP ratio: 16% of variance, F(1,64) = 251, p < 0.0001) and the type of TARP (%APC+ of GFP+: 12% of variance, F(7,64) = 24, p < 0.0001; APC/GFP ratio: 5% of variance, F(7,64) = 11, p < 0.0001). The individual flow data were fit to distinguish possible effects of threshold and slope, as in Figures 1B and 2A (Figure 3B) and further analyzed using a similar 3-way ANOVA on the homomeric data. GluA1, both flip and flop, showed greater thresholds than GluA2 while the heteromers were intermediate (Figure 3B). Indeed, 77% of the variance in the threshold values was accounted for by the subunit identity (F(1,64) = 874, p < 0.0001). Interestingly, we found that across TARPs, the flop subunits of GluA1 had lower thresholds than the flip variants, while the converse was true for GluA2 (Figure 3B, Figures S3C, S4C, Table 3, Subunit x splice variant interaction effect F(1,64) = 116, p < 0.0001).

**Figure 3.**
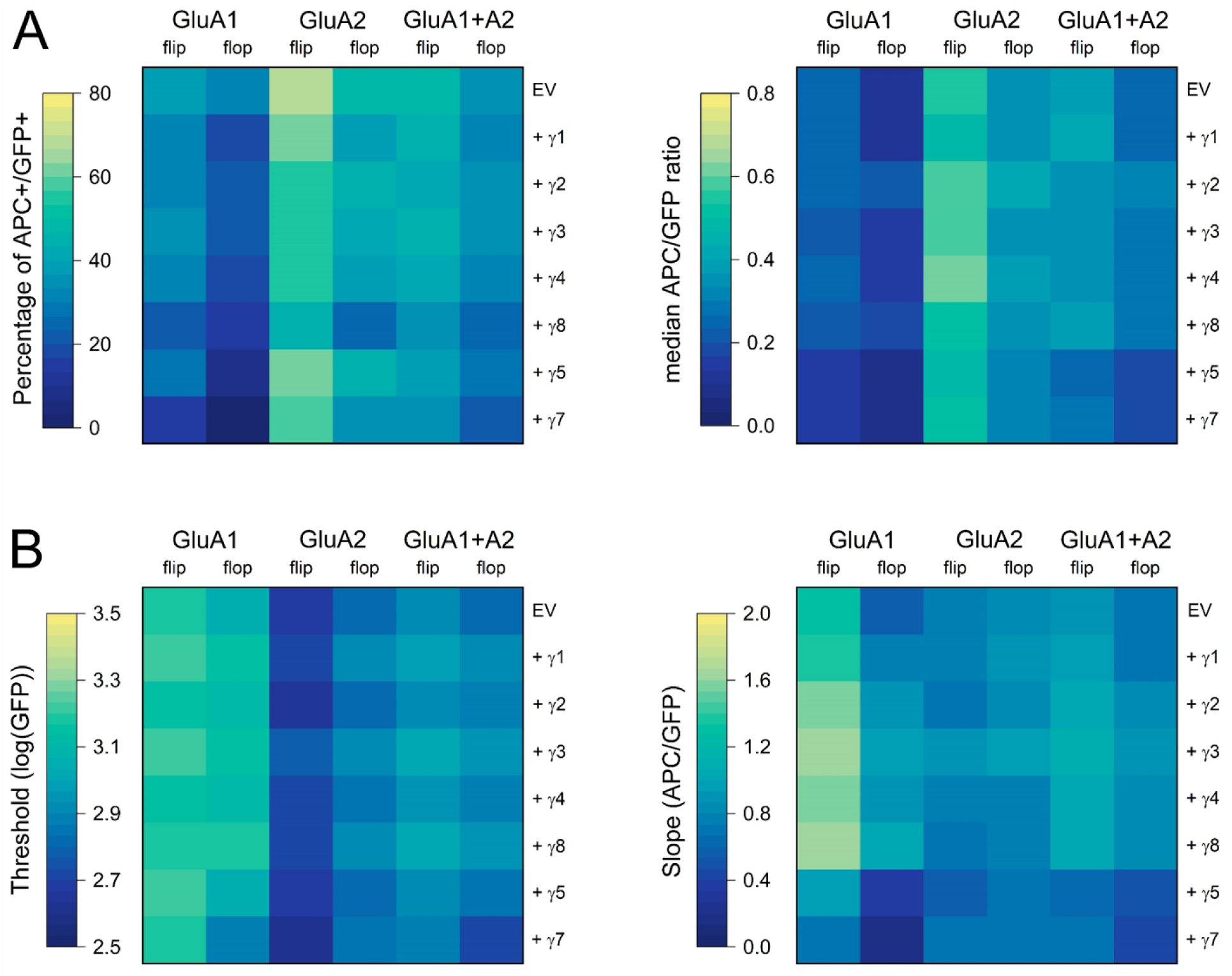
TARP effects on GluA1, GluA2 and heteromeric surface trafficking. **(A, *left)*** Heatmaps of percentage APC positive cells (APC+) out of GFP positive (GFP+) cells across all flow experiments for the indicated AMPA subunits and TARPs. EV stands for empty vector. (A, right) Heatmap of median APC/GFP ratio for all flow runs with the indicated AMPA subunits and co-transfected TARPs. (B) Heatmaps of threshold (left) and slope (right) fits for flow runs from GluA1, GluA2, or heteromers co-transfected with the indicated construct.

**Table 1:** Summary of percent jointly APC+ and GFP+ events.

|  | GluA1 |  | GluA2 |  | GluA1/GluA2 |  |
| --- | --- | --- | --- | --- | --- | --- |
|  | flip | flop | flip | flop | flip | flop |
| EV | 38.9<br>(1.7) | 22.4<br>(1.6) | 58.5<br>(8.7) | 47.7<br>(2.2) | 51.2<br>(7.1) | 38.6<br>(3.7) |
| $\gamma 1$ | 29.7<br>(3.9) | 17.0<br>(1.9) | 52.2<br>(4.3) | 39.9<br>(3.3) | 48.3<br>(7.2) | 31.4<br>(3.5) |
| $\gamma 2$ | 30.5<br>(2.0) | 20.2<br>(2.0) | 54.9<br>(5.5) | 44.3<br>(3.9) | 42.9<br>(4.4) | 36.9<br>(3.1) |
| $\gamma 3$ | 34.6<br>(2.5) | 22.7<br>(2.4) | 55.1<br>(7.9) | 43.0<br>(5.8) | 44.7<br>(5.5) | 35.6<br>(4.9) |
| $\gamma 4$ | 32.6<br>(2.4) | 18.7<br>(2.4) | 54.1<br>(4.8) | 39.5<br>(5.8) | 40.1<br>(3.7) | 32.3<br>(4.8) |
| $\gamma 8$ | 22.7<br>(1.5) | 13.9<br>(1.3) | 41.2<br>(5.4) | 25.5<br>(4.7) | 33.6<br>(4.5) | 23.5<br>(2.0) |
| $\gamma 5$ | 28.4<br>(3.4) | 9.5<br>(0.9) | 55.1<br>(4.1) | 43.8<br>(6.9) | 39.1<br>(5.1) | 29.1<br>(2.9) |
| $\gamma 7$ | 14.6<br>(2.5) | 2.2<br>(1.1) | 52.9<br>(4.3) | 34.7<br>(4.0) | 27.8<br>(11.3) | 16.3<br>(8.5) |

Data are means (standard deviation). N ranges from 3 to 8.

**Table 2:** Summary of median APC/GFP ratios.

|  | GluA1 |  | GluA2 |  | GluA1/GluA2 |  |
| --- | --- | --- | --- | --- | --- | --- |
|  | flip | flop | flip | flop | flip | flop |
| EV | 0.22<br>(0.01) | 0.12<br>(0.02) | 0.54<br>(0.13) | 0.36<br>(0.03) | 0.33<br>(0.08) | 0.25<br>(0.03) |
| $\gamma 1$ | 0.20<br>(0.02) | 0.13<br>(0.03) | 0.50<br>(0.04) | 0.35<br>(0.01) | 0.36<br>(0.06) | 0.25<br>(0.03) |
| $\gamma 2$ | 0.27<br>(0.02) | 0.20<br>(0.03) | 0.59<br>(0.09) | 0.40<br>(0.01) | 0.35<br>(0.07) | 0.29<br>(0.05) |
| $\gamma 3$ | 0.23<br>(0.01) | 0.17<br>(0.02) | 0.58<br>(0.09) | 0.37<br>(0.03) | 0.35<br>(0.09) | 0.26<br>(0.03) |
| $\gamma 4$ | 0.24<br>(0.03) | 0.15<br>(0.02) | 0.61<br>(0.04) | 0.38<br>(0.01) | 0.36<br>(0.09) | 0.27<br>(0.04) |
| $\gamma 8$ | 0.23<br>(0.01) | 0.19<br>(0.02) | 0.49<br>(0.06) | 0.33<br>(0.01) | 0.35<br>(0.08) | 0.26<br>(0.04) |
| $\gamma 5$ | 0.14<br>(0.01) | 0.09<br>(0.01) | 0.48<br>(0.02) | 0.30<br>(0.02) | 0.24<br>(0.05) | 0.20<br>(0.02) |
| $\gamma 7$ | 0.15<br>(0.01) | 0.10<br>(0.01) | 0.52<br>(0.03) | 0.31<br>(0.02) | 0.27<br>(0.09) | 0.18<br>(0.03) |

Data are means of median surface/total ratios from each flow experiment (standard deviation).
N ranges from 3 to 8.

**Table 3:** Summary of GFP thresholds.

|  | GluA1 |  | GluA2 |  | GluA1/GluA2 |  |
| --- | --- | --- | --- | --- | --- | --- |
|  | flip | flop | flip | flop | flip | flop |
| EV | 3.22<br>(0.05) | 3.05<br>(0.03) | 2.70<br>(0.05) | 2.83<br>(0.02) | 2.92<br>(0.13) | 2.81<br>(0.09) |
| $\gamma 1$ | 3.24<br>(0.06) | 3.15<br>(0.07) | 2.78<br>(0.08) | 2.92<br>(0.02) | 3.01<br>(0.14) | 2.90<br>(0.08) |
| $\gamma 2$ | 3.16<br>(0.07) | 3.13<br>(0.06) | 2.64<br>(0.02) | 2.82<br>(0.01) | 2.92<br>(0.10) | 2.85<br>(0.08) |
| $\gamma 3$ | 3.23<br>(0.07) | 3.14<br>(0.05) | 2.78<br>(0.07) | 2.91<br>(0.02) | 2.95<br>(0.14) | 2.90<br>(0.08) |
| $\gamma 4$ | 3.16<br>(0.03) | 3.12<br>(0.06) | 2.72<br>(0.05) | 2.86<br>(0.03) | 2.92<br>(0.11) | 2.87<br>(0.07) |
| $\gamma 8$ | 3.19<br>(0.07) | 3.18<br>(0.08) | 2.75<br>(0.05) | 2.91<br>(0.02) | 2.97<br>(0.14) | 2.93<br>(0.08) |
| $\gamma 5$ | 3.22<br>(0.06) | 3.04<br>(0.03) | 2.72<br>(0.04) | 2.82<br>(0.03) | 2.91<br>(0.12) | 2.80<br>(0.08) |
| $\gamma 7$ | 3.18<br>(0.08) | 2.89<br>(0.17) | 2.68<br>(0.03) | 2.84<br>(0.02) | 2.84<br>(0.14) | 2.87<br>(0.29) |

Data are means of log(GFP) channel signal intensity (standard deviation). N ranges from 3 to 8.

**Table 4:**
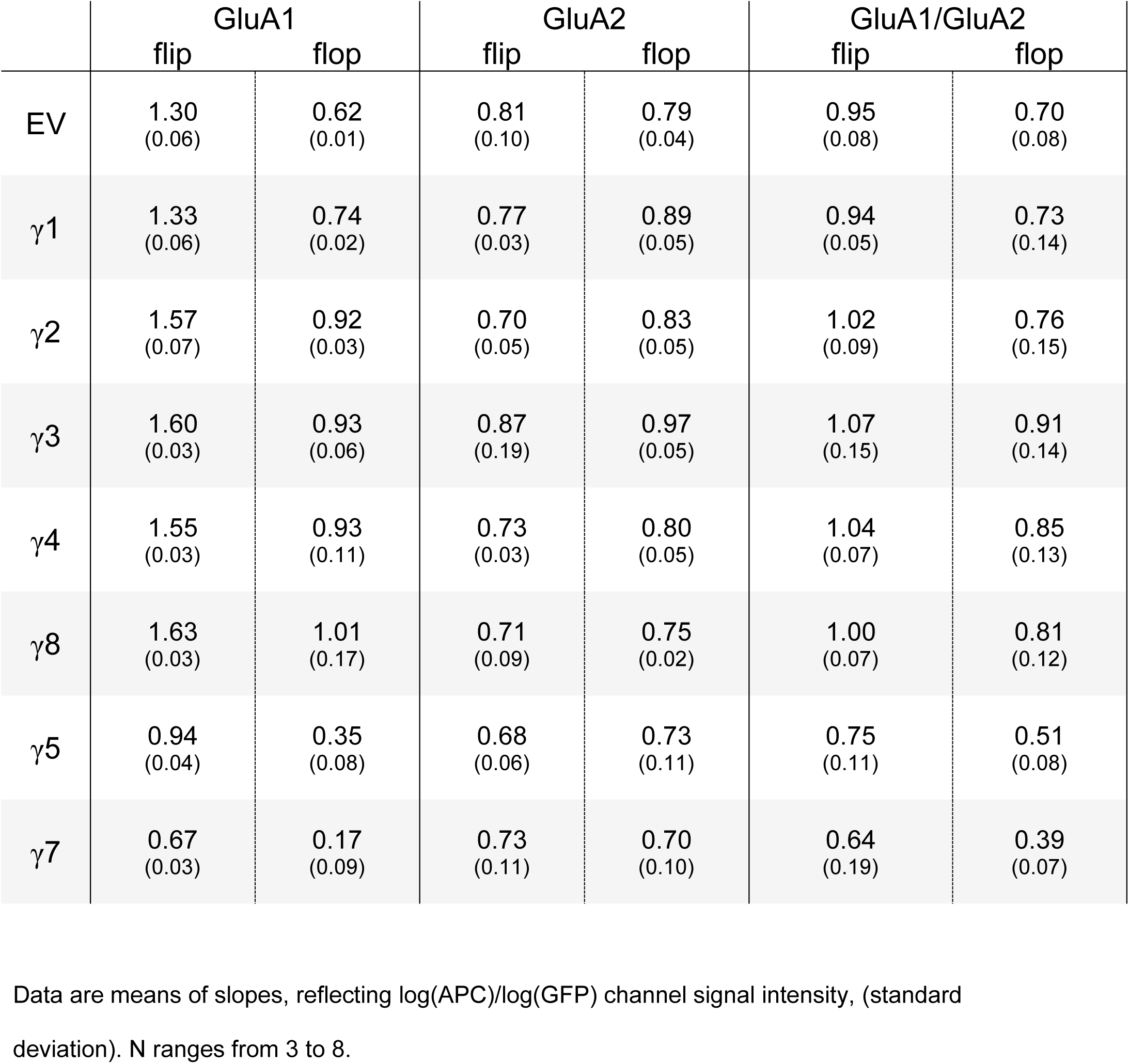
Summary of slopes.

Next, we examined the slope relating the total AMPA receptors to surface AMPA content (Figure 3B, right, Figures S3D, S4D and Table 4). Unlike prior measurements where either subunit identity or splice variant accounted for the majority of the effects in the data set, multiple factors and interactions between factors appear to influence the slope. The clearest effects were observed in the GluA1 flip, which possesses the largest baseline slope (Figure 3B, right, Figures S3D, S4D, and Table 3). Co-transfection of any type 1 TARP increased the slope of GluA1 flip while type 2 TARPs reduced it. A similar effect was observed with GluA1 flop, albeit with a lower intrinsic slope (Figure 3B, right, Figures S3D, S4D and Table 3). Interestingly, the slope of GluA2, both flip and flop, was not markedly altered by either type 1 or type 2 TARPs while that of heteromeric AMPAs was intermediate.

### TARP effects on AMPA trafficking assessed by surface luminescence

The flow cytometry-based assay separates trafficking effects into threshold versus slope at the cost of complexity and throughput. In addition, introducing fluorescent proteins at the carboxy terminus of AMPA receptors may compromise trafficking signals associated with the terminal PDZ ligand or intracellular domain. To complement the flow assay, we implemented a split luciferase-based approach. We appended an HA tag followed by an 11-amino acid segment of luciferase called HiBiT sequence immediately after the signal peptide and before the mature protein. HiBiT binds to the complementary large-bit protein (LgBiT) with sub-nanomolar affinity to form a functional nanoluciferase^45^. Measuring luminescence before and after cell permeabilization using digitonin yields surface and total measurements of AMPA receptors^46^ (Figure 4A). Using this approach, we measured the relative surface expression of GluA1 and GluA2, both flip and flop, as well as co-transfections of all possible flip/flop pairings (Figure 4B, left). We assessed the homomeric data using a 3-way ANOVA as above and, consistent with the flow data, found that subunit identity was the major driver of surface trafficking (47% of variance), with TARP and flip/flop cassette being secondary (22% and 17% of variance, respectively). Figure 4B summarizes these data as a heatmap of raw surface expression. To better visualize the effect of TARPs, we normalized the surface expression obtained with each TARP co-expression to the same-day surface expression of that AMPA subunit and splice variant co-transfected with an empty vector (Figure 4C). Across all AMPA subunits and splice variants, we found that type 1 TARPs increase surface expression while type 2 TARPs reduce it. Specifically, the fold change of type 1 TARPs versus empty vector ranged from 1.4 fold increase for GluA1 flop with γ3 to 2.3 fold greater with GluA1 flip and γ2 while the reduction by γ5 and γ7 ranged from 0.95 with GluA1 flop plus γ7 to 0.3 with GluA1 flip γ7 (Figure 4C). These data are further summarized in Figures S5 and S6 as well as Table 5.

**Figure 4.**
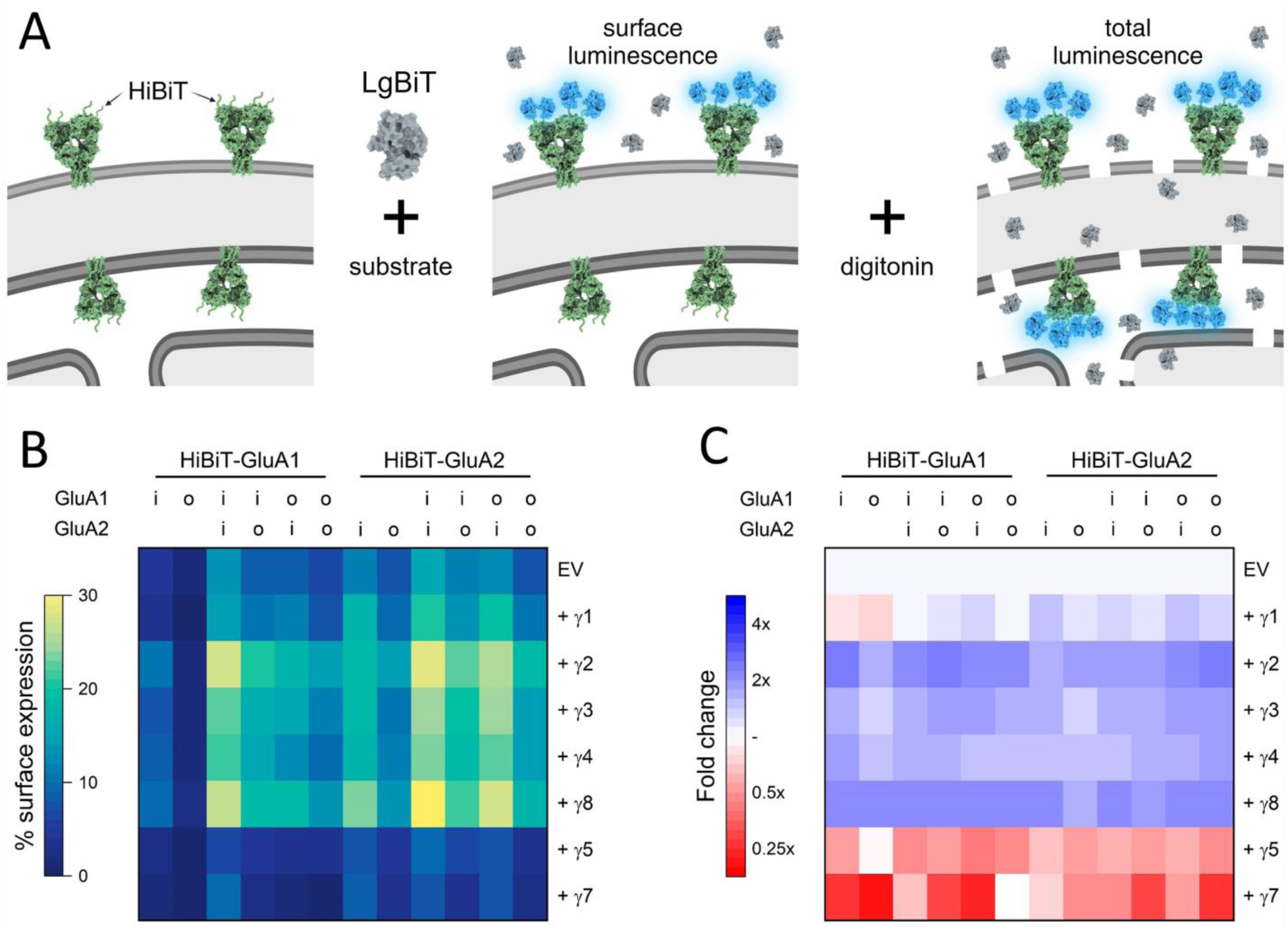
Homo and heteromeric GluA1 and GluA2 surface expression modulated by TARPs. **(A)** Schematic of HiBiT/LgBiT surface expression assay where an amino-terminal HiBiT tag binds with recombinant LgBiT to form functional nanoluciferase. Subsequent permeabilization by digitonin allows the separation of the surface AMPA from the total AMPA content. **(B,** *left)* Heatmap showing the percentage of surface expressed AMPA receptors for GluA1 and GluA2 homomers or GluA1/A2 co­transfection with either empty vector (EV) or the indicated TARP co-transfected. Flip and flop splice variants are denoted by *i* and o, respectively. **(B,** *right)* Heatmap of fold change in surface expression induced by co-transfection versus EV on the same day.

**Table 5:**
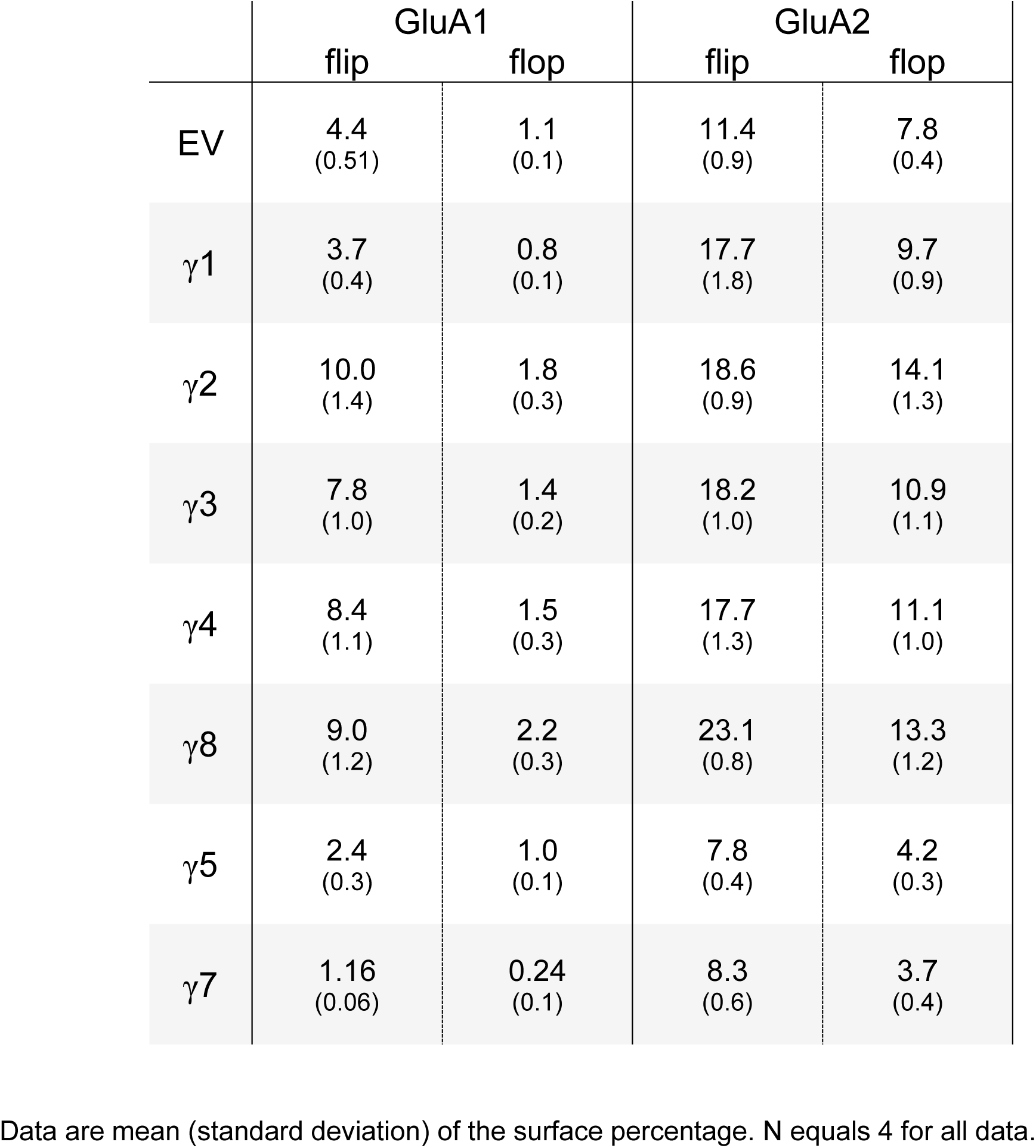
Percentage of homomeric AMPA receptors on cell surface based on luminescence.

|  | GluA1 |  | GluA2 |  |
| --- | --- | --- | --- | --- |
|  | flip | flop | flip | flop |
| EV | 4.4<br>(0.51) | 1.1<br>(0.1) | 11.4<br>(0.9) | 7.8<br>(0.4) |
| $\gamma 1$ | 3.7<br>(0.4) | 0.8<br>(0.1) | 17.7<br>(1.8) | 9.7<br>(0.9) |
| $\gamma 2$ | 10.0<br>(1.4) | 1.8<br>(0.3) | 18.6<br>(0.9) | 14.1<br>(1.3) |
| $\gamma 3$ | 7.8<br>(1.0) | 1.4<br>(0.2) | 18.2<br>(1.0) | 10.9<br>(1.1) |
| $\gamma 4$ | 8.4<br>(1.1) | 1.5<br>(0.3) | 17.7<br>(1.3) | 11.1<br>(1.0) |
| $\gamma 8$ | 9.0<br>(1.2) | 2.2<br>(0.3) | 23.1<br>(0.8) | 13.3<br>(1.2) |
| $\gamma 5$ | 2.4<br>(0.3) | 1.0<br>(0.1) | 7.8<br>(0.4) | 4.2<br>(0.3) |
| $\gamma 7$ | 1.16<br>(0.06) | 0.24<br>(0.1) | 8.3<br>(0.6) | 3.7<br>(0.4) |

Data are mean (standard deviation) of the surface percentage. N equals 4 for all data.

**Table 6:** Percentage of heteromeric AMPA receptors on cell surface based on luminescence.

|  | HiBiT-GluA1 |  |  |  | HiBiT-GluA2 |  |  |  |
| --- | --- | --- | --- | --- | --- | --- | --- | --- |
|  | A1i,<br>A2i | A1i,<br>A2o | A1o,<br>A2i | A1o,<br>A2o | A1i,<br>A2i | A1i,<br>A2o | A1o,<br>A2i | A1o,<br>A2o |
| EV | 13.0<br>(0.6) | 8.9<br>(0.5) | 8.6<br>(0.6) | 6.8<br>(0.5) | 15.6<br>(1.0) | 12.8<br>(1.0) | 11.6<br>(1.0) | 7.9<br>(0.9) |
| $\gamma 1$ | 14.0<br>(0.8) | 10.1<br>(0.7) | 11.1<br>(0.6) | 7.3<br>(0.6) | 20.5<br>(1.9) | 19.3<br>(1.9) | 13.9<br>(1.2) | 10.8<br>(1.6) |
| $\gamma 2$ | 27.1<br>(0.6) | 20.3<br>(0.6) | 17.2<br>(0.6) | 14.2<br>(0.4) | 28.9<br>(2.2) | 26.0<br>(2.0) | 22.5<br>(1.9) | 18.7<br>(2.0) |
| $\gamma 3$ | 22.5<br>(0.8) | 16.8<br>(0.7) | 15.2<br>(0.5) | 11.0<br>(0.4) | 24.9<br>(1.8) | 24.7<br>(1.9) | 19.8<br>(1.9) | 14.1<br>(1.7) |
| $\gamma 4$ | 21.1<br>(0.9) | 15.6<br>(0.3) | 12.8<br>(0.6) | 9.6<br>(0.3) | 23.4<br>(1.5) | 22.2<br>(1.4) | 18.1<br>(0.8) | 14.3<br>(1.4) |
| $\gamma 8$ | 26.5<br>(1.0) | 18.6<br>(0.5) | 18.1<br>(0.3) | 13.9<br>(0.6) | 31.4<br>(2.7) | 27.0<br>(2.2) | 21.7<br>(2.3) | 17.6<br>(1.9) |
| $\gamma 5$ | 6.3<br>(0.3) | 4.5<br>(0.2) | 3.6<br>(0.3) | 3.2<br>(0.2) | 9.0<br>(0.8) | 7.5<br>(0.5) | 6.0<br>(0.6) | 3.9<br>(1.0) |
| $\gamma 7$ | 4.2<br>(0.4) | 2.8<br>(0.3) | 2.0<br>(0.3) | 1.0<br>(0.5) | 7.4<br>(0.6) | 7.1<br>(0.6) | 3.5<br>(1.2) | 2.6<br>(1.3) |

Data are mean (standard deviation) of the surface percentage. N equals 4 for all data.

To assess the trafficking of heteromeric AMPA receptors, we repeated co-transfection experiments with the HiBiT tag on either the GluA1 or the GluA2 subunit. As expected, type 1 TARPs increased surface expression while type 2 TARPs reduced surface expression (Figure 4B and C, Figures S5 and S6, type 1 increases ranging from 1.4 fold to 2.4 fold; type 2 decreases ranging from 0.68 to 0.14 versus empty vector). Interestingly, we found that γ2 and γ8 increased surface expression of co-transfected GluA1 and GluA2 more than γ3 or γ4, respectively (Figure 4B and C, Supplemental S5 and S6). And we found that γ1 systematically elevated the surface expression of GluA2 but not GluA1 (Figure 4B and C, Figures S5 and S6). Given the effect on surface expression, we hypothesized that γ1 may act as a TARP with GluA2. However, co-transfection of γ1 with GluA2 did not increase the relative efficacy of kainate nor change the desensitization kinetics in excised patches (data not shown).

Finally, cell lines such as suspension HEKs may release transiently transfected proteins into the media through exosomes. Thus, some fraction of our surface expression signal may arise from these release proteins and/or dead cells. To control against a differential effect from such release, cells were split into two pools. One pool was washed with PBS prior to luminescence readings to remove released exosome proteins and/or dead cells while the other was not washed. These pools showed indistinguishable surface expression levels, indicating washing is not essential (Figure S7).

## Discussion

Using both flow cytometry and split nanoluciferase-based assays, we report the first systematic assessment of how TARPs, subunit type, and alternative splicing at the flip/flop cassette govern surface trafficking of AMPA receptors. We find that subunit identity is the major driver of surface expression, with GluA2 expressing much better than GluA1. Flow cytometry data analysis reveals this effect is primarily due to a lower threshold for channels to be delivered to the plasma membrane (Figures 1 and 3). Alternative splicing plays a secondary role, with flip subunits trafficking more effectively across the board than their flop counterparts in identical conditions (Figures 1, 3, and 4). Finally, the presence of TARPs influences surface expression, with type 1 TARPs (γ2-4 and γ8) enhancing surface expression while type 2 TARPs (γ5 and γ7) reduce surface expression (Figures 2, 3, and 4). These effects are conserved in co-transfection of GluA1 and GluA2, independent of which subunit is being detected (Figure 4B). We anticipate these assays will be useful in assessing the effects of post-translational modifications, ER resident chaperone proteins^8,47^ as well as patient mutations in both AMPA subunits^48,49^ and auxiliary proteins^50^. To the best of our knowledge, no other study has systematically assessed trafficking across as many conditions. However, multiple other studies have examined the effects of either GluA1 versus GluA2^43^, flip versus flop^12,43,51,52^ or a subset of TARPs on specific AMPA receptors^14,31,53,54^. Our work aligns with these past investigations in that type 1 TARPs enhance the surface expression of all AMPA subunits, regardless of flip or flop. This contrasts functional modulation by specific TARPs, where γ2 strongly affects the desensitization kinetics of GluA2 flip but not on GluA2 flop^55^. We also observe that γ5 reduces surface expression, as reported previously^54^, although this effect was not detected in another study^31^. And our observation that γ7 is a negative regulator of surface trafficking aligns with the increase in AMPA currents from granule cells upon γ7 knockdown^56^.

### Limitations of current work

All our measurements are conducted in HEK suspension cells. These cells differ from neurons in several key respects. HEK cells lack the complement of ER-resident chaperone proteins, auxiliary proteins, and endo/exocytosis machinery present in neurons (although see ^13,44^). Beyond differences in protein complement, neurons may utilize both canonical and non-canonical secretory pathways for somatic versus dendritically synthesized proteins^7^. Nor can HEK cells accurately reflect the influence of neuron-specific glycosylation on these receptors^57^. In addition to these shared limitations, the flow cytometry assay also requires that a fluorescent protein be appended to the channel. We added the fluorescent protein to the C terminal tail, potentially interfering with PDZ or other motifs. However, since our flow and luminescence data strongly agree, adding the C terminal fluorescent protein does not appear to be a major confound. Despite the limitations, HEK suspension cells offer considerable advantages. These include ease of use, reproducibility, and the ability to scale. Also, because suspension cells lack endogenous AMPA receptors and TARP proteins, it is possible to quantify the effect of one specific TARP on one specific AMPA receptor without having the genetically delete or compensate for all the other subunits as one would need to do in neurons. Thus, these assays are well suited to the study of *de novo* disease-associated variants in either AMPA receptors or auxiliary/chaperone proteins in isolation.

### Source of intrinsic differences between subunits

GluA2 shows more abundant surface expression than GluA1 (Figure 1, 4B), consistent with past work^43^. Flow cytometry analysis reveals GluA1 has a higher “threshold” than GluA2 (Figure 1E), likely accounting for the difference in surface expression. We suggest this greater “threshold” arises due to a longer maturation time of GluA1. The molecular basis for this longer maturation time is unclear but may reflect differences in glycosylation requirements. GluA1 possesses 6 N-glycosylation sites, two of which (N63 and N363) are critical for HEK cell surface expression^58^. In contrast, GluA2 has four N-glycosylation sites, with only one of these (N370) being essential for surface expression in HEK cells^59^. The more stringent glycosylation requirement of GluA1 may lead to intracellular accumulation and account for the greater threshold prior to surface expression (Figure 1E). Another possibility is that GluA1 homomers are more difficult to fold properly in HEK cells, leading to increased misfolded proteins accumulating in the ER. In this view, the “threshold” measurement we observe does not reflect GluA1 homomers awaiting surface delivery but an accumulation of misfolded protein. Future work using pulse chase assays, domain swapping and mutations will be able to determine the molecular basis for this subunit difference.

Within the last decade, many disease-associated variants in GRIA genes have been reported^1,49,60^. These variants may possess alterations in channel gating, trafficking, or both^1,49^. In addition, new chaperone proteins have been uncovered, shedding light on the intricate biogenesis of AMPA receptors^8,13,20,47^. The throughput of methods developed here are ideally suited to assay trafficking across variants and with specific chaperone proteins. Thus, despite decades of research on AMPA receptors, new technologies and approaches continue to offer up new questions and avenues of inquiry.

## Acknowledgements

We thank the University of Rochester Medical Center’s Flow Cytometry Resource, especially Steven Polter, for instrument training and troubleshooting.

## Author Contributions

T.C. and T.W.M. conceived of the flow cytometry and split nanoluciferase portions of the project, respectively. T.C. and T.W.M. designed and performed experiments analyzed results and interpreted data. D.M.M analyzed data, interpreted results, and drafted the manuscript. All authors contributed to the editing of the manuscript and approved the final version.

---

**Figure S1.**
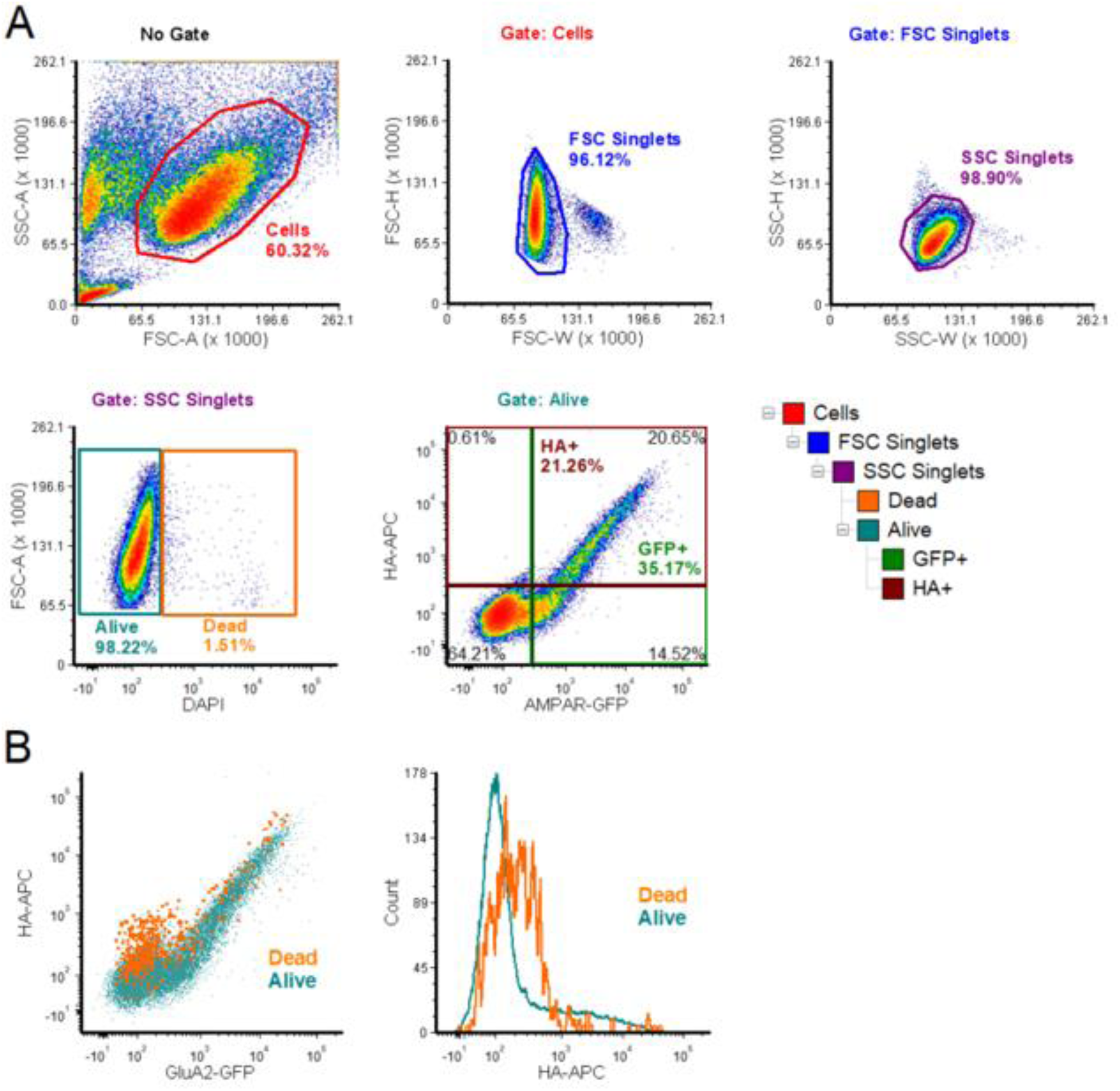
Gating scheme with DAPI stain to remove false positive ‘surface’ signal. **(A)** Gating scheme with forward and side scatter (FSC and SSC, respectively) gates as well as DAPI positive exclusion gate and quadrant plot. **(B)** Example GFP versus APC scatter (*left*) and APC histogram (*right*). DAPI positive cells (labelled Dead) are shown in orange and excluded cells in teal.

**Figure S2.**
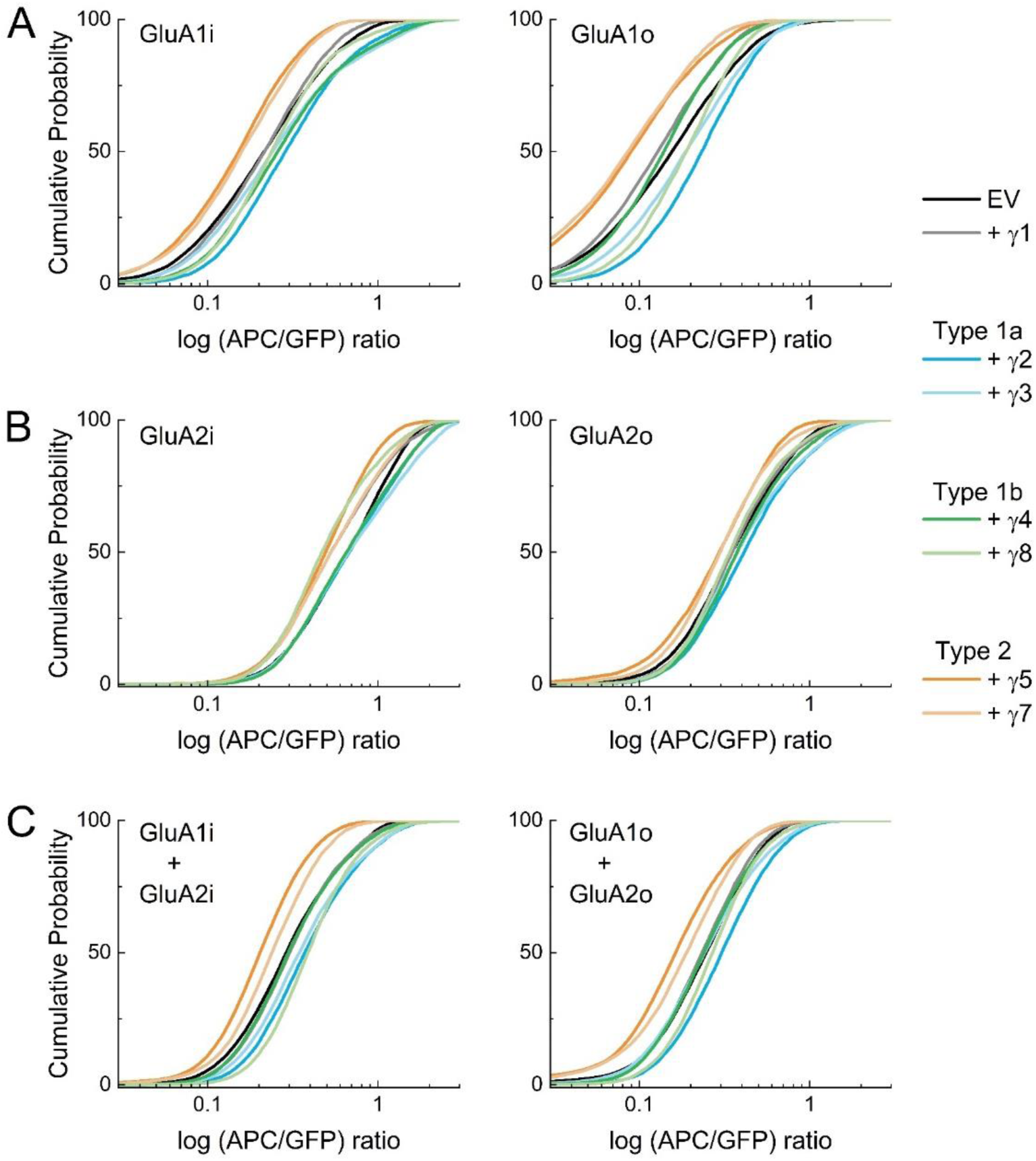
Type 1 TARPs enhance while Type 2 TARPs impair surface trafficking of GluA1 and 2, both flip and flop. **(A-C)** Single cell APC/GFP ratio cumulative probability plots for GluA1 **(A)**, GluA2 **(B)** and GluA1+GluA2 **(C)**, both flip (*left*) and flop (*right*) variants co-transfected with the indicated gamma subunits.

**Figure S3.**
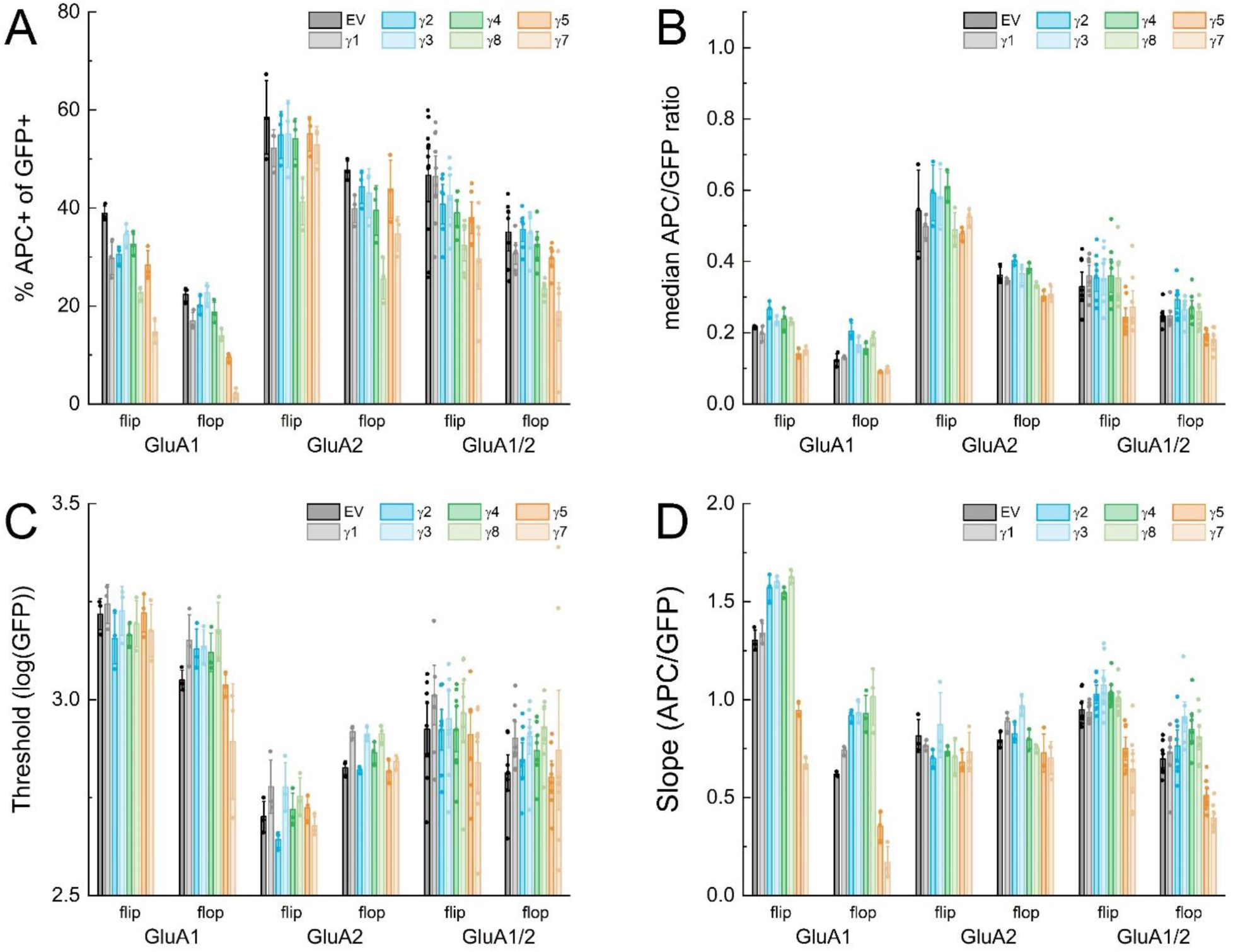
Type I and 2 TARPs have opposing positive and negative effects on surface trafficking of GluA1 and 2, both flip and flop. **(A, B)** Summary plots of percentage APC positive events of GFP positive events **(A)** and single cell APC/GFP ratios **(B)**. Symbols show the median value from a single flow experiment, and column height, and error bars show the mean and SEM across flow experiments. **(C, D)** Summary of piecewise linear fit thresholds **(C)** and slopes **(D)** across flow experiments. Symbols show the fit value from individual flow experiments, while column height and errors bars depict the mean and SEM across experiments.

**Figure S4.**
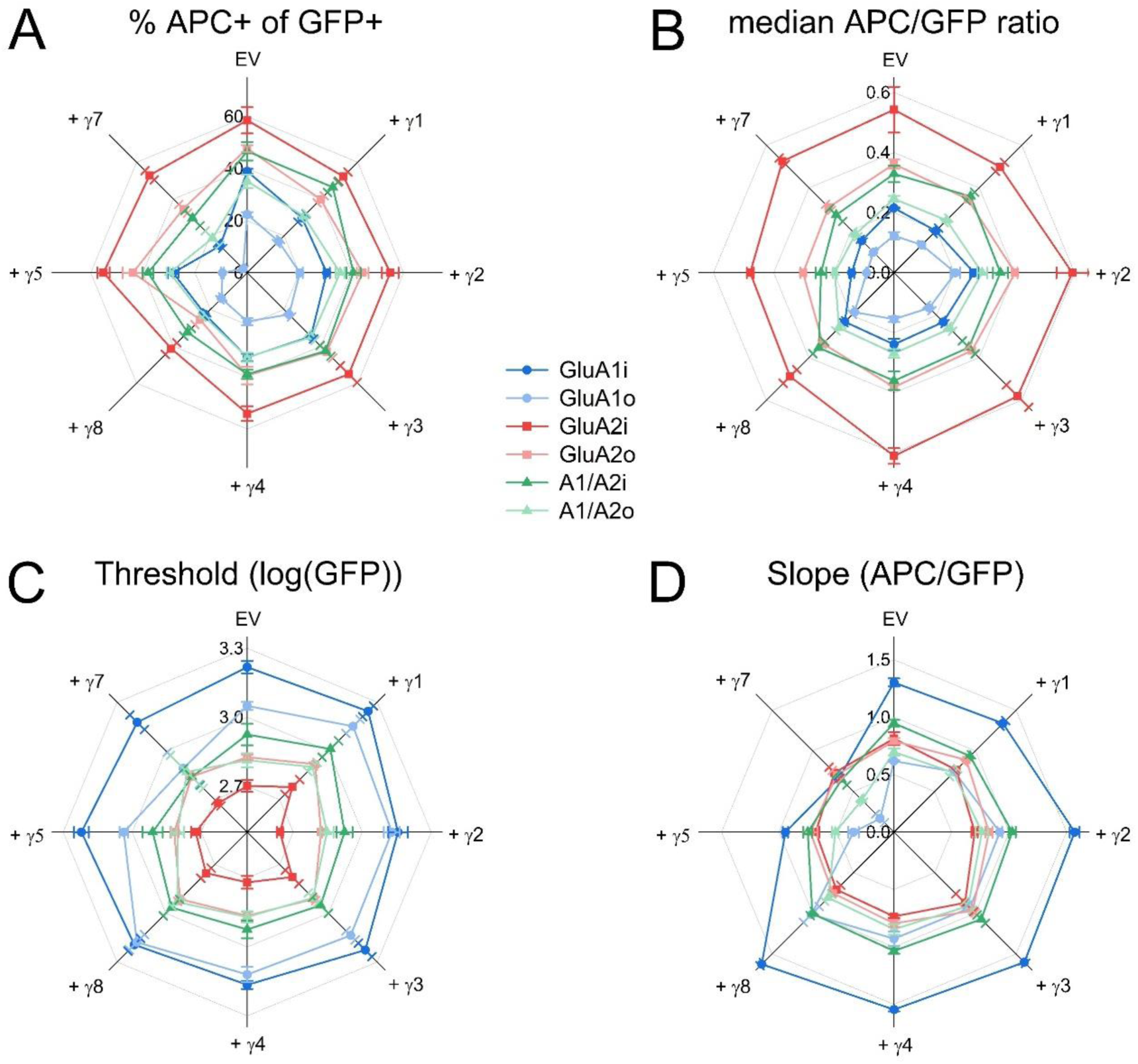
Type I and 2 TARPs have opposing positive and negative effects on the surface trafficking of GluA1 and 2, both flip and flop. **(A, B)** Radar plots of percentage APC positive events of GFP positive events **(A)** and median APC/GFP single cell ratios **(B)**. **(C, D)** Summary radar plots of thresholds **(C)** and slopes **(D)** across flow experiments. Symbols and error bars show the mean and SEM, respectively, across flow experiments.

**Figure S5.**
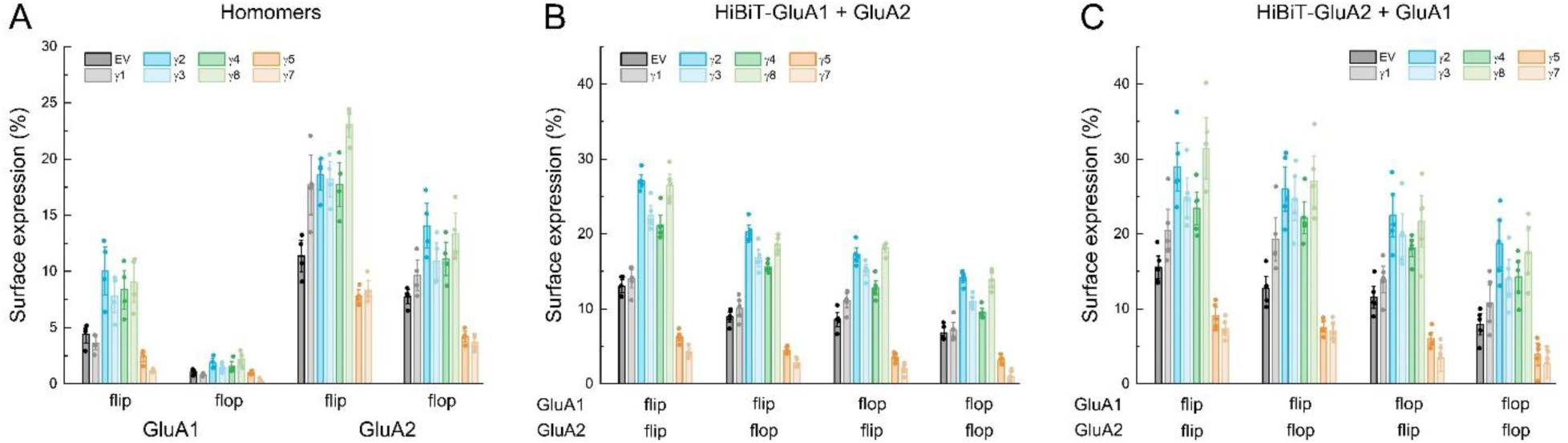
TARPs impact both homomeric and heteromeric surface expression. **(A)** Summary plot of AMPA surface expression for GluA1 or GluA2, either flip or flop variant, when co-transfected with the individual TARP. **(B)** Summary plot of GluA1 surface expression when co-transfected with GluA2 and the indicated TARP. **(C)** Same as in **B** but for GluA2. Symbols show individual experiments and error bars are SEM.

**Figure S6.**
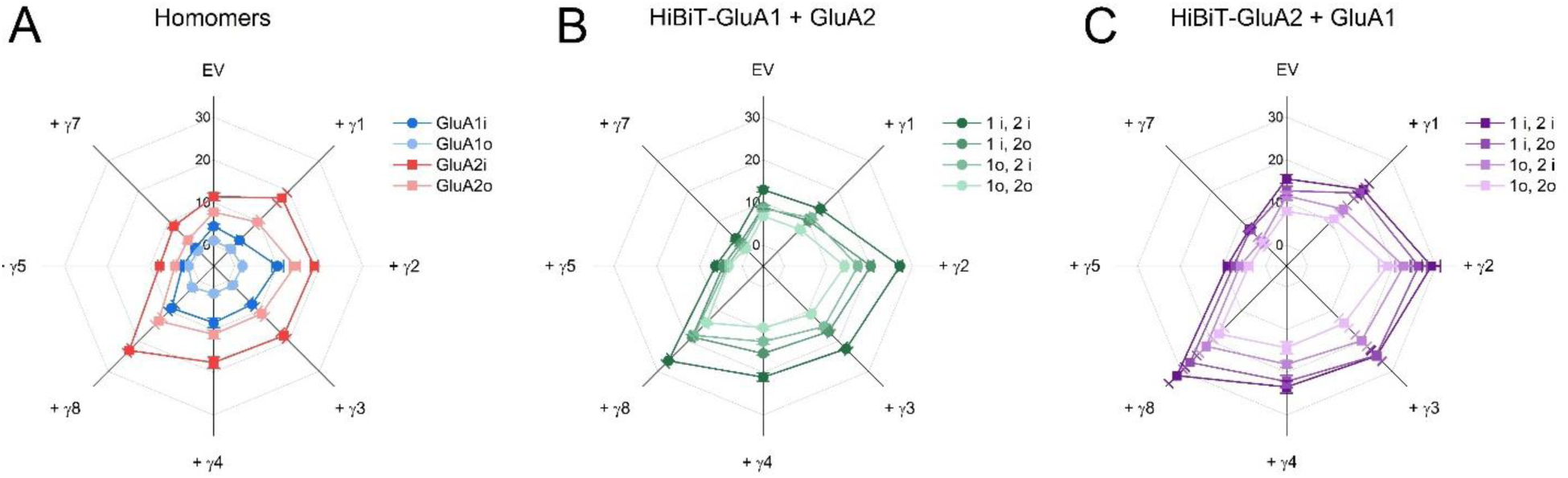
Radar plots of TARP impact on homomeric and heteromeric surface expression. **(A)** Radar plot of AMPA surface expression for GluA1 or GluA2, either flip or flop variant, when co-transfected with the individual TARP. **(B)** Summary plot of GluA1 surface expression when co-transfected with GluA2 and the indicated TARP. **(C)** Same as in **B** but for GluA2. Symbols show mean of independent experiments and error bars are SEM. Flip and flop are represented by *i* and *o*, respectively.

**Figure S7.**
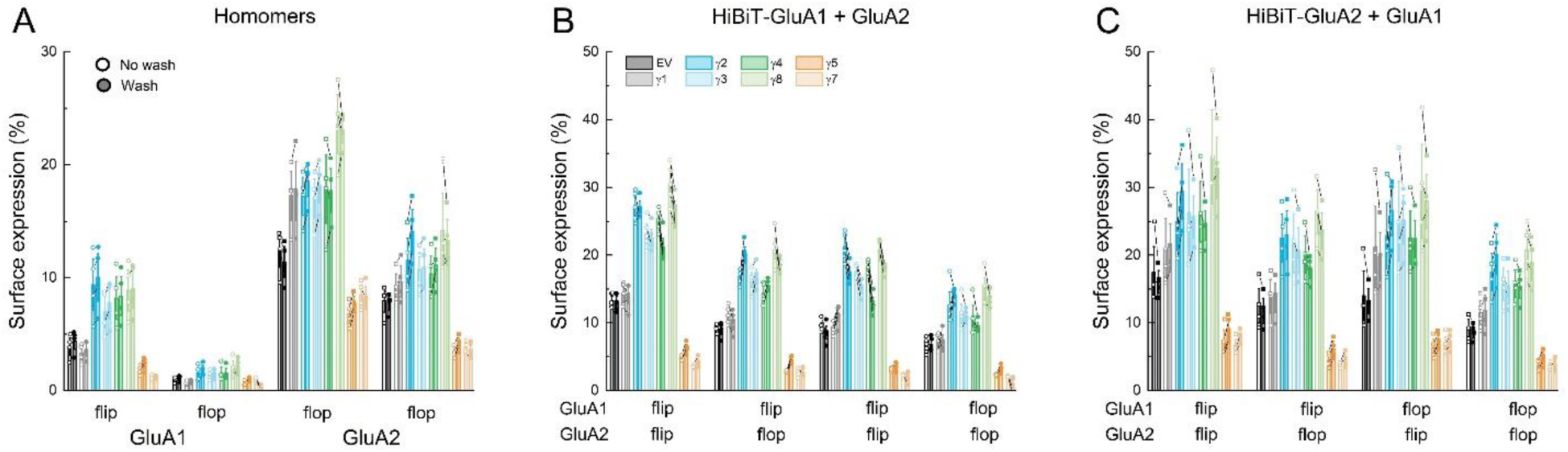
Washing has negligible impact on surface expression results. **(A-C)** Summary of surface expression for homomeric GluA1 or GluA2 **(A)**, or GluA1 **(B)** or GluA2 **(C)** when co-transfecting both subunits. Open symbols and bars are results from cells directly from growth media. Filled symbols are following wash steps. Symbols show individual experiments connected by lines and error bars are SEM.

